# Characterisation of *Saccharomyces cerevisiae* and *Saccharomyces jurei* hybrids for oenological applications

**DOI:** 10.64898/2026.10.30.763130

**Authors:** William Royle, Kohsheen Pandita, Rakshitha Rangarajan, Silvia Carlin, Francesco Reale, Alistair Hanak, Urška Vrhovšek, Daniela Delneri

**Affiliations:** Manchester Institute of Biotechnology, University of Manchester, M1 7DN Manchester, UK; Division of Evolution and Genomic Sciences, School of Biological Sciences, Faculty of Biology Medicine and Health, University of Manchester, M13 9PT Manchester, UK; Foundation Edmund Mach, Via Edmund Mach 1, 38010, San Michele all’Adige, TN, Italy

## Abstract

The *Saccharomyces* genus present as a genetically diverse set of microorganisms capable of tolerating a range of environmental stressors, demonstrating high fermentation capacity and producing interesting flavour compounds. Such diversity can be exploited in a single organism by producing interspecies hybrids, overcoming the limitations of a single species. Here, as proof of principle, we exploited a library of F12 interspecies diploid hybrids between *Saccharomyces cerevisiae* and *Saccharomyces jurei* (Sc/Sj), to identify new strains harbouring traits of oenological relevance, such as ethanol and glycerol production, superior tolerance to acetic acid, limited production of H_2_S, and optimal growth on fructose. Moreover, we calculated robustness across strains tested on solid media comparing the trade-off between performance on a single conditions and overall robustness. Progeny from a range of robustness values with favourable H_2_S production profiles were isolated for further analysis via micro and small-scale fermentations. Micro-scale fermentations in synthetic grape must containing individual wine-associated stressors were performed, and the robustness of the selected strains was confirmed. The wine characteristics of the hybrids were tested in small-scale fermentations in synthetic grape media. We found that both ethanol and glycerol production across our selected progeny matched or exceeding that from the commercial wine strain, and most hybrids produced less acetic acid and more fatty-acids, including higher quantities of associated esters. Overall, we were able to identify suitable Sc/Sj hybrids with interesting aroma profiles to develop further for wine fermentation.

## Introduction

Yeasts are an increasingly commercially relevant species in a range of biotechnologies including wine production, beer production and industrial ethanol production from a variety of feedstocks [1]. Each application presents its own challenges, environmental stressors impact the growth, production efficiency, and in the case of beverage production, flavour profile [2, 3].

Wine production accounts for a large proportion of global revenue associated with microbial fermentations [4]. While traditional processes are highly valued there remains a market for development of wines with new flavour profiles and characteristics. The yeast used in the production of wine are not only crucial to the final alcohol percentage, but also significantly influence the flavour of the final product [5].

Yeast of numerous species are known to exist in initial grape must, with various species involved in spontaneous fermentation [6-8]. However, *Saccharomyces* species generally outcompete other species present and go on to dominate the fermentation [8]. *Saccharomyces cerevisiae* is crucial for the primary fermentation producing ethanol from the glycolysis of glucose and fructose present in grape must. Commercial strains are favoured for their ability to withstand the harsh environment of grape must and ability to effectively ferment [9, 10].

Numerous strategies exist to develop unique flavours and concepts for wine and beer production. There is an interest in the use of native *Saccharomyces cerevisiae* strains to provide distinct organoleptic qualities linked to the grape variety, geographical location and production methods, termed the *Terroir* [11, 12]. Mixed fermentations are also valued, non-saccharomyces species can provide greater complexity and also contribute to control of spoilage yeasts [13-15] and performing controlled mixed fermentations is an attractive strategy for development of unique wines [16].

Within the *Saccharomyces* genus itself, there are a range of interesting phenotypes, crucially temperature tolerance differs between species. *Saccharomyces eubayanus*, *jurei* and *kudriavzevii* are more cold-tolerant species [17-20]. For example, *Saccharomyces jurei* strains D5088 (NCYC3947) and D5096 (NCYC3962) display optimum growth at 27.8°C and 27.2°C, respectively [20].

Conducting wine fermentations at lower temperatures can lead to different qualities, including an increase in fruity aromas [21, 22], this has been associated with improved preservation of volatile compounds that constitute fermentative aroma [23].

Inter-species hybridisation is relatively common within the *Saccharomyces* genus and exists both in nature and anthropogenic environments [24, 25]. Indeed, the most famous example *Saccharomyces pastorianus* (syn. carlsbergensis), an allopolyploid hybrid of Saccharomyces cerevisiae and Saccharomyces eubayanus, arose in lager beer fermentation [18], and arguably constitutes the most valuable yeast hybrid being the most widely used in alcoholic beverage fermentations [26]. Lager beer is produced at lower temperatures (5-14°C) than the majority of ale type beers (15-20°C) [27], providing the necessary selection pressure for cold tolerance. *S. pastorianus* harbours favourable traits from both parents, a phenomenon termed heterosis or hybrid vigour, its cold tolerance from the *S. eubayanus* parent and superior fermentation kinetics from the *S. cerevisiae* lineage [28].

Hybridisation between saccharomyces species is also common in wine production, both occurring naturally [29] and through targeted approaches [30]. *Saccharomyces cerevisiae* and *S. uvarum* hybrids have been constructed to improve oenological function at lower temperatures [31].

Hybridisation offers vast prospects for the development of novel yeast strains, which can be exploited to improve fermentation and enhance the production of flavour compounds in specific environments, yielding unique, industrially relevant phenotypic traits [32, 33]. *Saccharomyces jurei*, the most recent addition to the *Saccharomyces* genus [34] remains relatively unexplored in terms of its biotechnological potential. Not only do the two strains identified in the French alps show improved cold tolerance compared to *S. cerevisiae* [20], but interesting metabolite profiles have been observed in fermentations with *S. jurei* strains. Here, higher production of medium chain fatty acid esters and relatively high concentrations of higher mass alcohols including isobutanol and iso-amyl alcohol are produced in comparison to *S. eubayanus* and *S. pastorianus,* as well as a notable phenolic clove like flavour note [35]. Of which increased production of 4-vinyl-gluanicol, phenolic compounds and fusel alcohols are likely to contribute [36]. Hybridisation with *Saccharomyces cerevisiae* enhances the fermentative potential of *S. jurei*, improving ethanol yield whilst maintaining the cold tolerant phenotype and interesting aroma profile [36].

Robustness, as a principle, provides a measure of a systems capability to maintain functionality across different perturbations [37]. Differing definitions are recorded, yet in biological systems, robustness can be considered as strain phenotype in response to restrictive or stressful conditions [38].

Recent work has identified a crucial need for the evaluation of robustness when developing biotechnological production strains [38, 39]. This has been applied conditions associated with industrial ethanol fermentation using lignocellulosic hydrolysate as the feedstock [40]. Presently, quantifying robustness has been neglected in the characterisation of strains for the production of wine, yet grape must is a complex matrix containing a mix of carbon sources, weak acids, antifungal agents present from cultivation of grapes and other stressors [41] and provides a unique set of stressors for fermentative yeast [2, 42, 43].

In previous work, we generated tetraploid hybrids between different strains of two species, namely *S. cerevisiae* and *S. jurei* (*Sc/Sj*), for which none of the parent strains is known to be involved in beverage fermentation [34, 44, 45]. Allopolyploidisation allows the tetraploids to undergo meiosis to produce diploid hybrid spores, which were randomly crossed and put through 12 meiotic cycles (F12) to allow ample recombination events [46].

Here, as proof of principle, we exploited genetically diverse fertile interspecific tetraploid hybrids between *S. cerevisiae* and *S. jurei,* and their progeny, to test the potential of *S. jurei* in winemaking, using a commercial *S. cerevisiae* wine strain for comparison (summarised in Figure 1). Collections of F12 diploid progeny were screened in stressor conditions associated with grape must and wine production specific traits including growth on fructose, acetic acid and SO_2_. Phenotypic data allowed the calculation of robustness, using methods described in Trivellin et al. 2002 [40]. Moreover, specific hybrid progeny exhibited an equal, or in some cases, lesser production of H_2_S in comparison to the commercial wine yeast strain. Furthermore, we investigate the growth kinetics of interesting progeny and the commercial strain in media mimicking grape must, we calculate robustness for different indicators of growth kinetics. By scaling up fermentations we can measure, via HPLC and GC-MS, the yields of favourable compounds; ethanol, glycerol and various volatile organic compounds (VOCs). We find no significant differences in ethanol yield between hybrids and the commercial strain, with specific hybrids producing more glycerol and greater quantities of desirable ethyl-esters.

**Figure 1.**
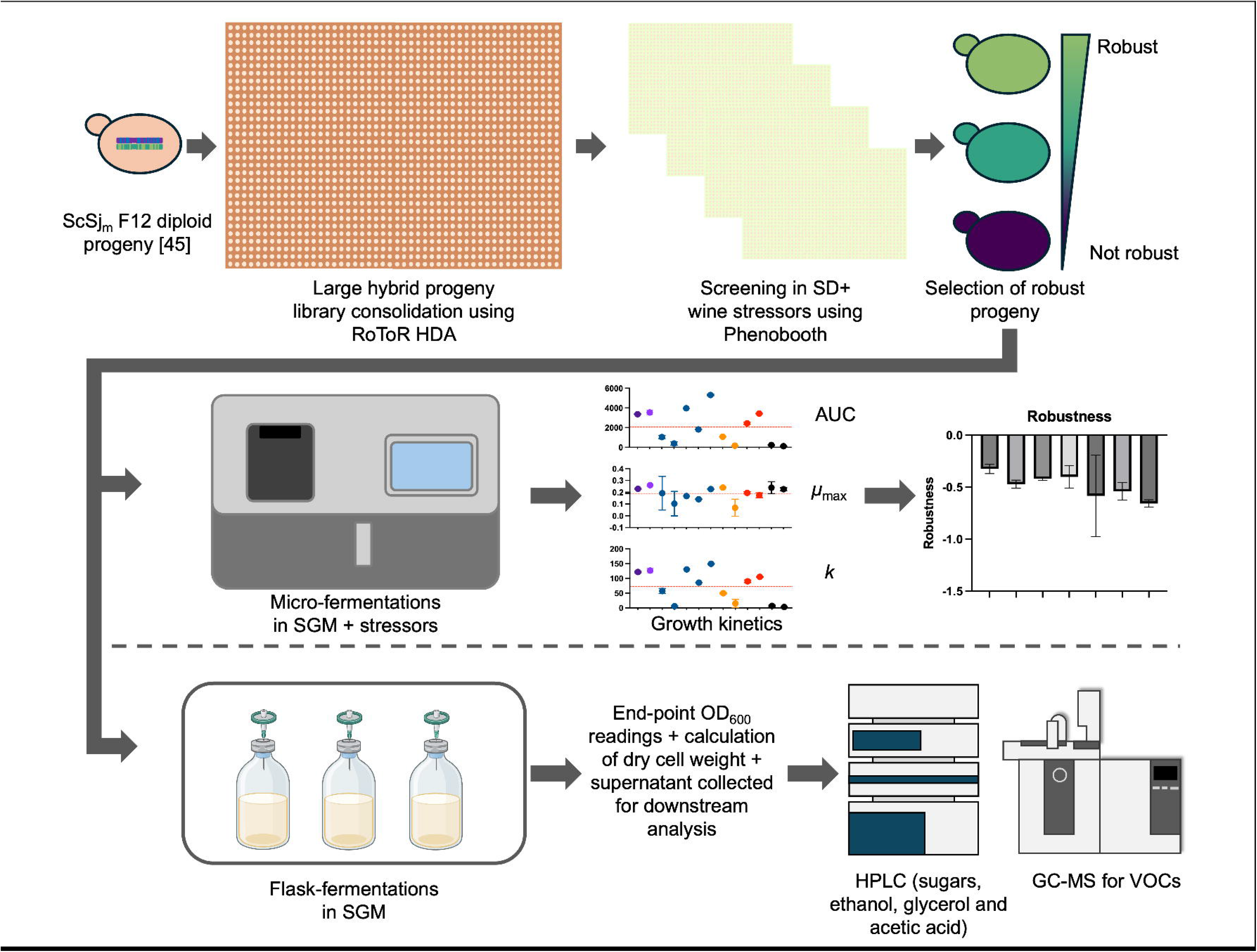
Experimental overview. Phenotypic screening on solid SD media containing different wine stressors (fructose, acetic acid, potassium metabisulphite and ethanol) allowed calculation of robustness using colony size as a proxy for spore fitness. Progeny with interesting characteristics were selected for higher resolution investigation into growth kinetics following micro-fermentations (Biolector) and calculation of robustness for AUC, maximum growth rate (*µ*_max_) and carrying capacity (*k*). Small-scale flask fermentations are also performed allowing calculation of key wine chemicals including aroma compounds.

Within this paper we test ScSj hybrid progeny generated from parent strains with no known background in fermented alcoholic beverage production; *S. jurei* strains D5088 and D5095, isolated from high altitude Oak tree bark (*Quercus*) [34], *S. cerevisiae* YPS128 (OS3) from north American oak and *S. cerevisiae* strain DBVPG6765 belongs to the Wine/European clade [44, 47] (although its specific origin is listed as unknown). Selecting robust progeny with good properties for wine production. Overall, this study underscores the great potential of these hybrids and widescale use of interspecific hybridisation for the generation of improved wine industry strains, generating novel and interesting aroma profiles.

## Results and Discussion

### Phenotypic screening of hybrid collection

Multigenerational diploid offspring of tetraploid hybrids [46] containing the genome of four strains belonging to two different species (Table 1), namely *S. cerevisiae* strain OS104 (*Sc^OS104^*) and OS3 (*Sc^OS3^*), and *S. jurei* strain D5088 (*Sj^D5088^*) and D5095 (*Sj^D5095^*) were used to investigate growth changes in the presence of acetic acid (0.3%, and 0.5%), fructose (2%), potassium metabisulfite (Kmet) (200mg/L and 500mg/L), and ethanol (14% and 15%), at two different temperatures (20°C and 30°C). High-throughput phenotyping on solid media [48] allowed us to assess the divergence of phenotypes within the F12 generation (Figure 2), and to identify spores with interesting growth characteristics.

**Figure 2.**
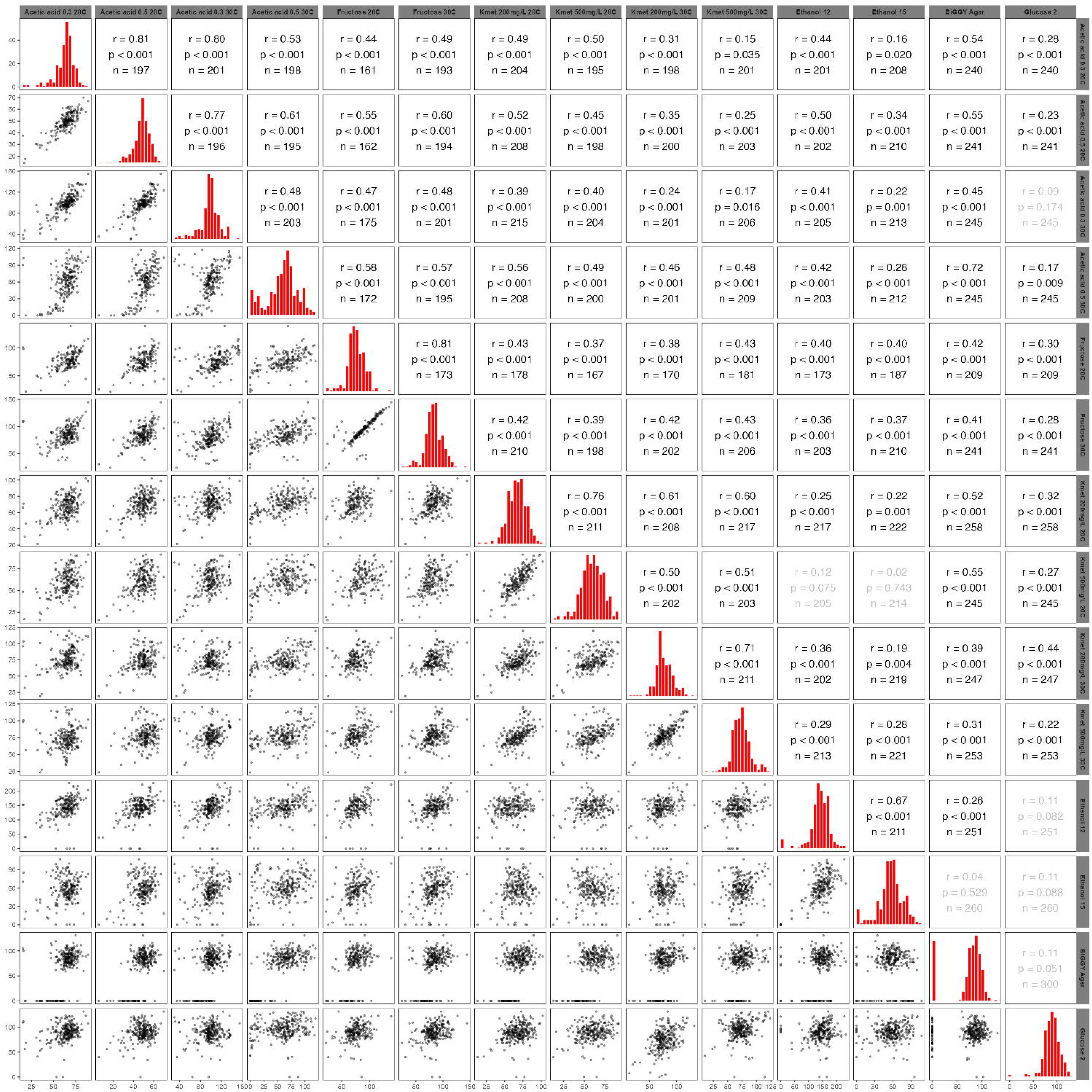
Scatter plot matrix displaying fitness (colony size) from high-throughput phenotypic screening on wine stressors. Phenotypic variance is shown for F12 hybrid collections of ScSj parental genomes [45]. Each dot on the scatter plots represents the median colony size (across four technical replicates) for a single F12 hybrid. Histogram plots show distribution of phenotypes across the population. Pearson correlation coefficients were calculated for each pair of phenotypes and *r* is displayed here for values with significance of *p* < 0.001(black text) or *p* >0.001 (grey text).

**Table 1.**
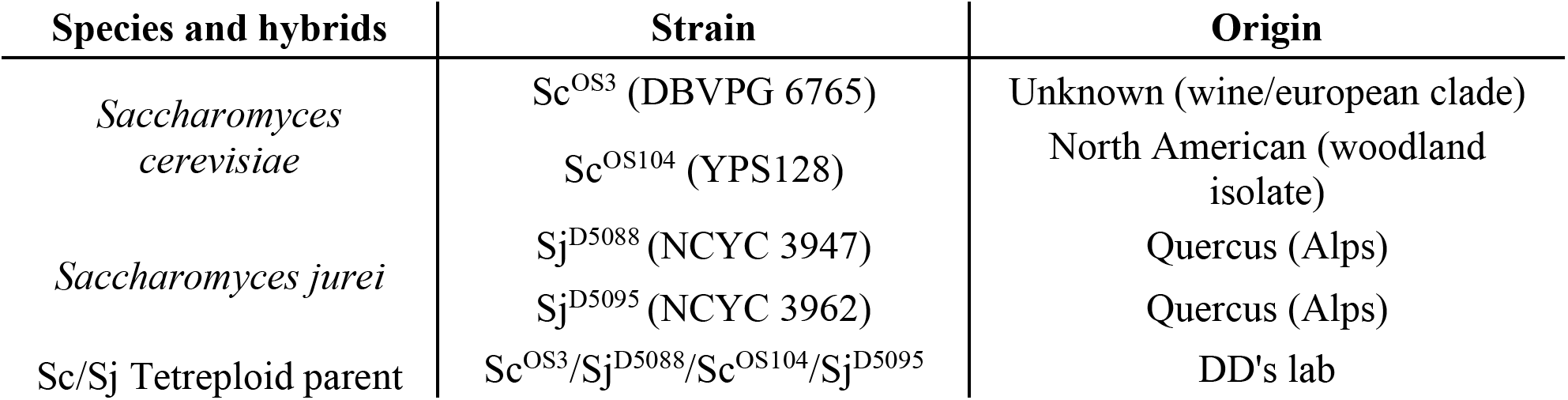
Background and origins of all parental strains and hybrid tetraploids used for phenotypic screening.

Across all conditions tested we see a normal distribution in phenotypes (diagonal histogram plots, Figure 2). As with previous screening of interspecies hybrid progeny, we observe a range in phenotypes across the population of hybrid progeny [46, 49], quantified as the coefficient of dispersion (COD) with a maximum COD of 0.28 in 0.5% acetic acid at 30°C condition (Table S1). In the majority conditions tested, we observe a positive correlation between perturbations, as presented by the Pearson correlation factor (Figure 2). Within conditions in which the same stressor was applied at different growth temperatures, we observe a broadly consistent response from spore to spore, notably in 2% fructose a Pearson correlation of 0.81 (P<0.0001) is observed. A positive correlation between conditions is also observed for acetic acid tolerance, across concentrations and temperatures tested (Figure 2), with the strongest linear correlation observed between 0.3% and 0.5% at 20°C (r=0.81). At higher temperatures a weak positive correlation is observed when the concentration of acetic acid is increased (r=0.48), suggesting a threshold for stress response in more extreme environments. We observe a slightly stronger positive correlation between Kmet and acetic acid at 20°C (r=0.39 to r=0.59) than at 30°C (r=0.24 to r=0.46). A Resistance to SO2 has been associated with resistance to carboxylic acids previously [50, 51]. And at low level concentrations of sulphite, the Haa1p regulon may confer some resistance to sulphite stress, which may account for the positive correlation observed here. However, in higher burden conditions the overlap in genes upregulated in both acetic acid and SO_2_ stress is more modest, suggesting somewhat separate resistance mechanisms [52].

Separately, we are aware of ScSj interspecies hybrids harbouring a number of pleiotropic genes governing fitness in acetic acid and growth in low temperature [46]. Low temperature was not tested in isolation in this study, but pleiotropic stress response genes may play a role in the positive linear correlation we observe between acetic acid conditions and low temperature potassium metabisulphite (Kmet) conditions (Figure 2). We also observe a weak positive relationship in 2% fructose at both temperatures and across all potassium metabisulphite conditions (200ug/mL and 500ug/mL).

To further examine the best performers per condition we performed spot assays, comparing strains from the upper and lower quartiles to an industrial wine strain and to the parental tetraploid. Of particular interest was the resistance to acetic acid, the top performers identified show greater tolerance to acetic acid at 0.3% and 0.5% than the commercial strain (Figure S1). The function of this screening was to select strains of interest for further study, as with previous studies [46, 49], we see that hybridisation can provide interesting characteristics, with extreme phenotypes possible.

### Colony screening confirms trade-offs in robustness and performance

We calculated robustness across solid media screening, using colony size as a proxy for fitness. We applied the method described in Trivellin et al. 2022 [40], using the fano factor to calculate an estimate for robustness (*R*) across all stressors tested (Figure 3, Figure 4A). Using this method a strain with high robustness has an R value close to 0 [40]. We did not include the score for H_2_S production (light intensity on Biggy Agar) within the calculation for robustness, to maintain data uniformity. We compared performance in each stressor with overall robustness (Figure 3). We found no strong positive linear relationship between performance (fitness) in each of the conditions tested, confirming previous work demonstrating trade-offs in robustness and performance [39] and supporting the general idea that robustness is inherent to trade-offs in specific performance [53] although it is worth noting these results pertain to a single phenotypic quality (colony size). However, a weak correlation (Pearson coefficient of 0.31 – 0.35) was observed in higher concentrations of acetic acid at both temperatures tested (Figure 3). Suggesting a strain with greater robustness performs better in weak acid stress.

**Figure 3.**
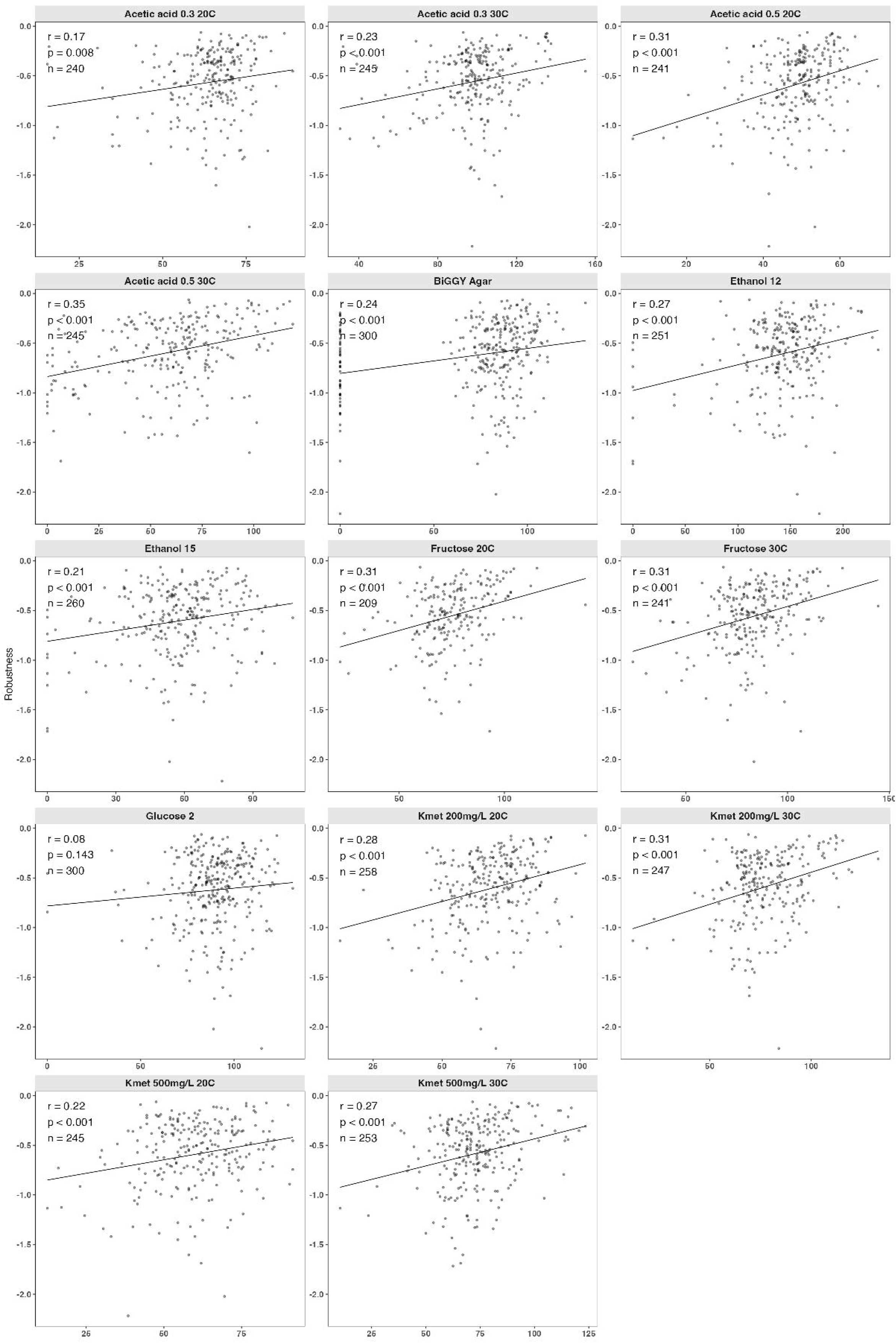
Comparison between phenotypes of F12 progeny in response to wine stressors vs robustness of the progeny. Robustness is calculated for each F12 hybrid using fitness data across all stressor conditions tested, formula is that used in Trivellin et al. 2022 [39]. Pearson correlation coefficient is calculated for each phenotype-robustness comparison with a significance of *p* > 0.001 for all comparisons except 0.3%(v/v) acetic acid 20°C and 2% glucose at 20°C.

**Figure 4.**
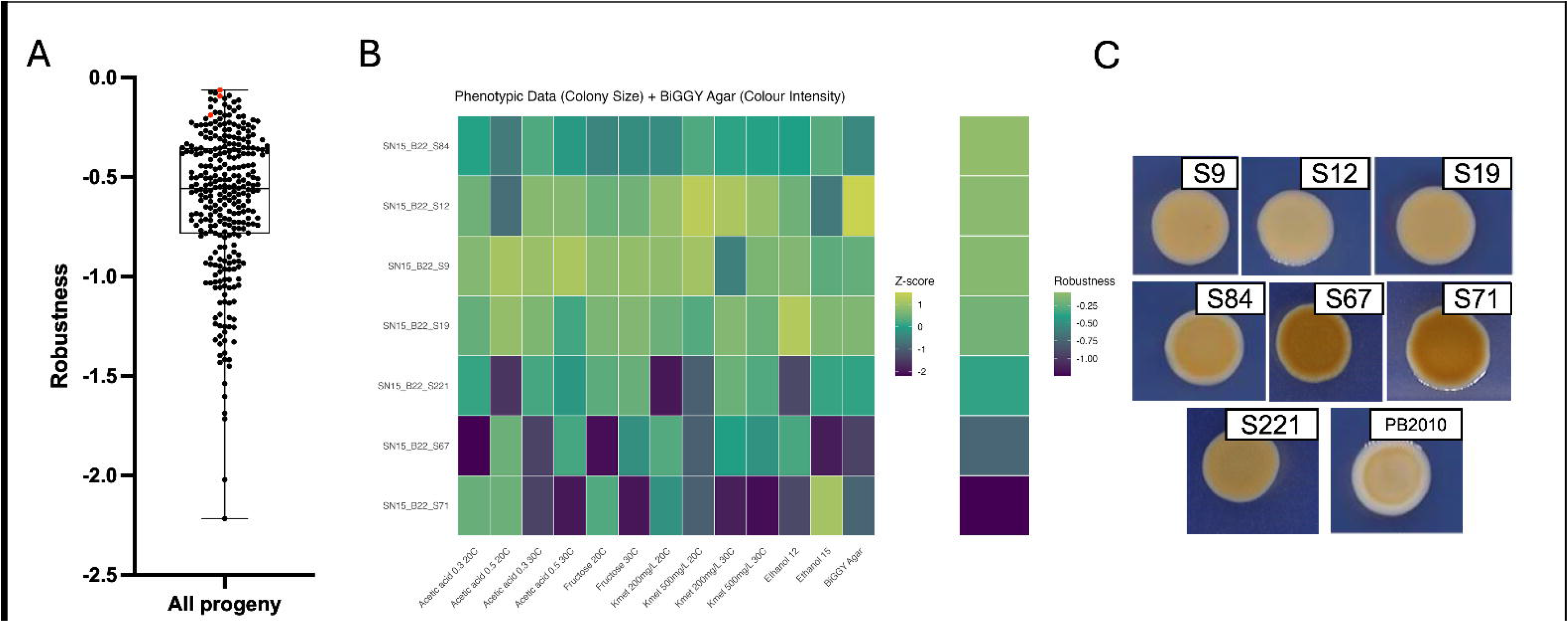
Selection of F12 hybrids with different levels of robustness for higher resolution investigation. **A** Box plot showing robustness, as calculated from colony size data, across the total ScSj F12 population, each dot representing a single F12 hybrid. Highlighted in red are hybrid progeny selected for micro- and small-scale fermentations in SGM. **B** Heatmap showing overall phenotype (fitness) of selected progeny in wine stressors, hybrid progeny are shown in order of robustness (secondary scale) top being more robust, bottom less robust. **C** Spot assay of selected hybrid progeny on BiGGY agar, darker coloured colonies show greater H_2_S production, lighter less H_2_S production. Selected hybrid progeny show a range in colour with S12 and PB2010 the lightest.

Acetic acid exists largely in its undissociated form at typical wine pH, therefore, can freely diffuse across the lipid bilayer leading to accumulation of acetate and an increase in intracellular pH. Weak organic acid (WOA) tolerance relies on a number of pathways in *Saccharomyces* including maintenance of cytosolic pH through proton efflux [54, 55], the cell wall integrity (CWI) pathway, the high-osmolality glycerol (HOG) pathway and the calcineurin pathway [56, 57]. Potential genetic advantages that exist in hybrid progeny to WOA stress may result in more general advantages to chemical stress and hence improve general robustness. One of the principal ideas behind biological robustness is system redundancy [37], such redundancy is intrinsic to the WOA stress response, which involves multiple pathways and systems for resistance [56, 57].

To pick out potential strains of interest, we isolated 7 strains from the overall collection with a range of robustness values (highlighted points, Figure 4A), and H2S production scores (light intensity). Screening data and robustness values are summarised in Figure 4B, S84 demonstrated the highest robustness from all progeny tested, with an overall R value of -0.06 the lowest robustness spore selected was -1.25 for S71. We performed spot assays in BiGGY agar to confirm high-throughput results for H_2_S production, summarised in Figure 4C. The darker the spot colour the grater the production of H_2_S, we see a range of production across selected spores with lightest colour from PB2010 and hybrid S12 (Figure 4C).

### Robustness analysis of selected progeny in liquid synthetic grape media

To further investigate selected progeny, we performed micro-scale fermentations (1.5mL total volume) utilising Biolector. Fermentations were performed in liquid SGM media without the addition of high concentration glucose and fructose, all conditions contained 2% (w/v) glucose, excluding the 2% (w/v) fructose condition. Thirteen stressor conditions typical of grape must were tested and scatter light readings collected at regular intervals. All growth data was analysed using GrowthCurveR. To evaluate robustness we utilised the method described in Trivellin et al. 2022 [40]. Robustness was calculated separately for each characteristic of growth dynamics, the area under curve (AUC) establishing a broad summary of growth, the maximum growth rate (*µ*_max_) and the maximum population size that can be sustained in a particular condition or carrying capacity (k).

An initial comparison of intrinsic growth rate across the hybrid progeny and PB2010 in 2% glucose shows a significantly higher growth rate for PB2010 (Figure 5A, Figure S2) with an average of 0.288hr^-1^ across replicates. The commercial strain also has a higher growth rate in 2% fructose (0.248hr^-1^). Demonstrating overall higher performance of the commercial strain in favourable conditions. Yet, when considering performance across multiple stressors, a number of hybrid strains demonstrate greater robustness for growth rate (*r*) (Figure 5B). Growth rate is of particular importance when considering an industrial fermentation strain, this can reduce the risk of potential spoilage yeasts from dominating the fermentation [58].

**Figure 5.**
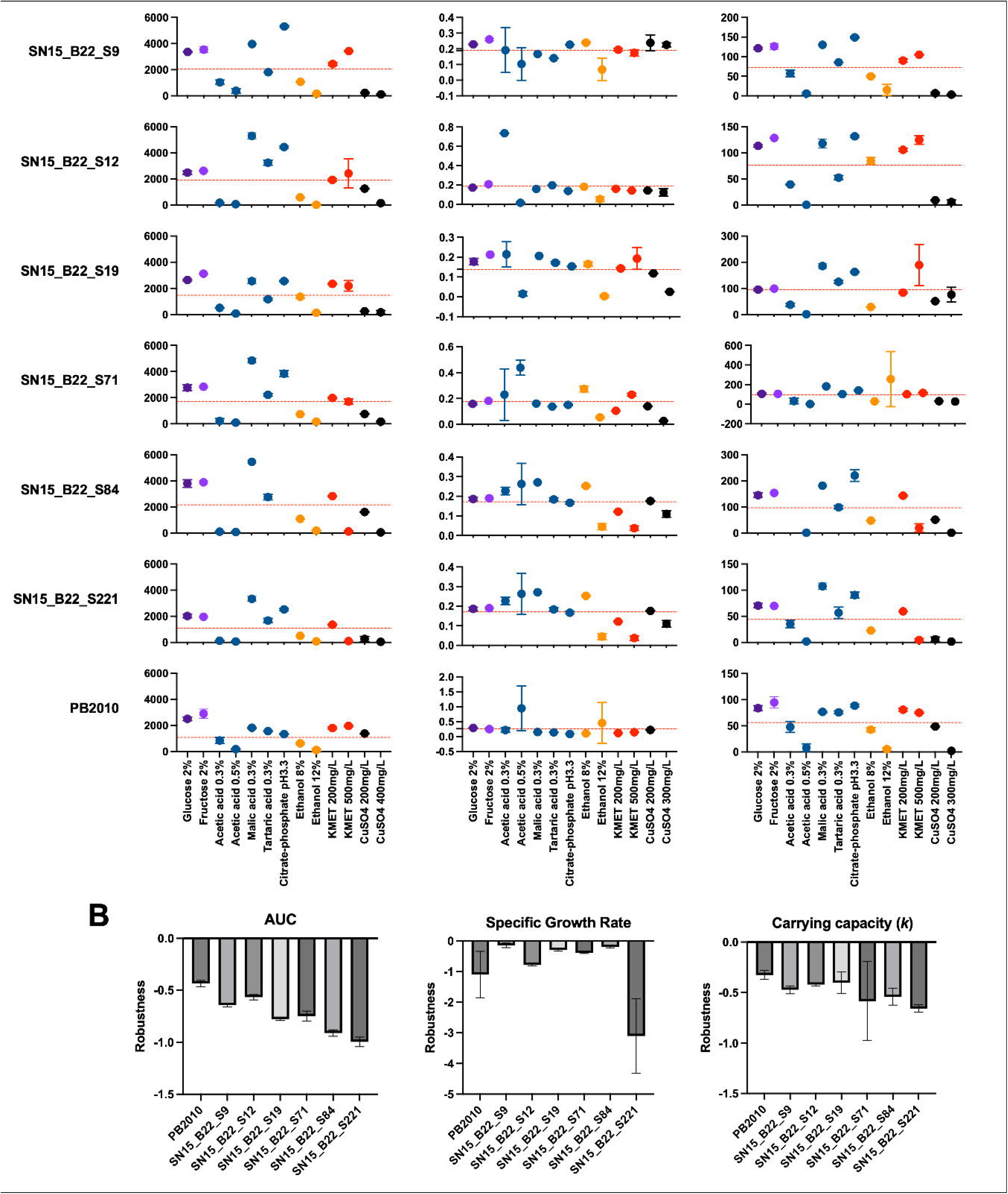
Selection of F12 hybrids with different levels of robustness for higher resolution investigation. **A** Growth kinetics for each of the hybrid progeny and for the commercial strain, following growth assays (Biolector micro-fermentations). Area under the curve (AUC) is calculated from the area under the logistic curve, accounting for growth rate, and the biomass. Max growth rate (*µ*_max_) and carrying capacity (*k*) give further indications of a strains ability to tolerate a condition. Each indication is calculated for the 13 conditions and presented here. **B** Robustness, as calculated using the fano factor method [39] across AUC, *µ*_max_, and *k*. The closer the R value to 0 the higher the robustness for that factor. PB2010 showers greater robustness when considering AUC, yet does is not as robust as hybrid progeny S9, S19 and S84 when considering *µ*_max._

Robustness was slightly greater for the industrial strain when considering AUC and the carrying capacity (Figure 5B), which is not surprising given the environment PB2010 has evolved to thrive in grape must fermentation. The difference in robustness rank observed between solid media screening in SD media and liquid micro-fermentations in modified-SGM (Table S2) is potentially due to the difference in media between the two screening methods, minimal SD media for solid phenotyping and liquid SGM for micro-fermentations.

Therefore, we suggest that media selection may have a greater impact on experiments to assess robustness, we propose that future experiments to resolve robustness characteristics must consider appropriate media for the specific application.

### Flask fermentations using select hybrids

Flask fermentations were performed to compare the general performance of selected hybrid progeny to a commercial wine strain. To establish clear and unambiguous relationships between the hybrid progeny examined and the commercial strain, PB2010, fermentations were performed in complete synthetic grape must (SGM), as described in Viana et al. 2014 [59] with 250g/L total sugars. End-point fermentation samples were analysed for sugar attenuation and glycerol, ethanol and acetic acid production by HPLC.

High sugar attenuation was ubiquitous across all fermentation samples, with no-detectable glucose or fructose in end-point samples. Ethanol concentration was matched across all strains, with no significant difference in yield (g ethanol/g initial sugar) as shown in Figure 6A. With yields of around 0.447 to 0.477 across all strains tested (111g/L to 119g/L) (Figure 6A, Figure S3, Table S3).

**Figure 6.**
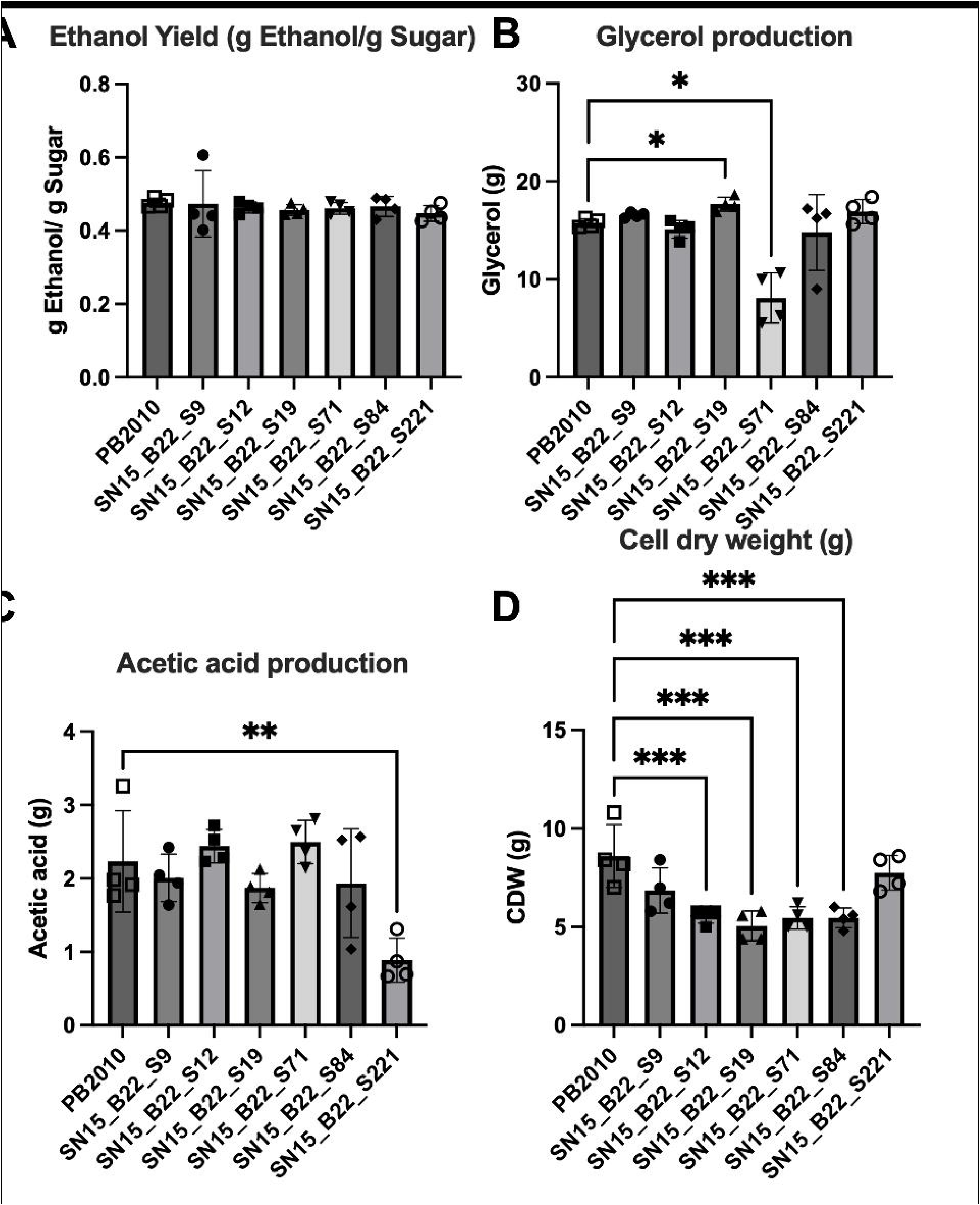
Key indicators of fermentation performance from flask fermentations in SGM. **A** Ethanol production was quantified via HPLC and yield (g Ethanol/g initial sugar) calculated. No significant differences were observed between hybrid strains and PB2010. **B and C** Glycerol production (g) quantified via HPLC in end-point fermentation samples and acetic acid production (g) also quantified via HPLC (see methods). **D** Cell dry weight (g) measured following 7 day fermentations in SGM. Significance across all data sets (**A**,**B**,**C** and **D**) was determined using ANOVA where; * indicate *p* < 0.05, ** indicate *p* < 0.005 and *** indicate *p* < 0.0005.

Hybrid strain S19 produced a more glycerol than PB2010, with final concentrations of 17.69g/L and 15.73g/L respectively (Figure 6B, Table S3). In yeast synthesis of glycerol serves two main functions in *Saccharomyces*, principally the synthesis serves to balance NADH/NAD redox during anaerobic fermentation [60]. Glycerol also acts as an osmoprotectant [61].

Glycerol does not contribute directly to aroma, yet may function in wine to improve overall organoleptic properties at concentrations of ≥ 10g/L [62]. Previous work has associated changes in glycerol content with the viscosity of a wine [62, 63] and this is widely accepted in the community [64]. However, it is not clear how perceptible glycerol concentration is to overall sensory characteristics of the beverage, with typical wine concentrations ranging from 4g/L to 9g/L [65]. And artificially increased concentrations of glycerol added to batches of wine, to a final concentration of 10.2g/L, had mixed results in sensory testing [66].

Greater interest is afforded to the potential to shift metabolic flux away from ethanol production, potentially towards glycerol production for the generation of less lower ethanol wines [67]. However, this can lead to undesirable consequences such as increased production of acetic acid and acetoin, volatile compounds associated with poorer quality wines [65, 68, 69]. Engineering strategies now exist that offer improved production of glycerol and reduction in ethanol generation without overproduction of acetate or acetoin [70-73].

The majority of hybrid progeny showed no observable difference in acetic acid production when compared with the commercial strain. S221 demonstrated a significantly lower production of acetic acid than PB2010 (Figure 6C), with concentrations of 0.89g/L and 2.23g/L, respectively. Acetic acid concentrations in wine typically range from 0.1g/L to 0.5g/L in wines [74], the concentrations we observe here are higher than would be seen in natural grape juice fermentations. This is likely a result of the composition of the SGM used in fermentations, however, optimisation of SGM media is beyond the scope of this study. Grape must factors influencing acetic acid production are well defined and may be explained by nitrogen and amino acid deficiencies [75], glucose content or due to inadequate concentrations of long chain unsaturated fatty acids [76, 77].

Cell-dry weight was calculated to provide an estimate of final biomass. Hybrid strains S12, S19, S71 and S84 show significantly lower final biomass than the commercial strain (Figure 6D), with average measurements ranging from 5.05g to 5.60g in hybrids and 8.60g for PB2010.

### Interspecies hybrids produce more fatty-acids and associated ethyl-esters

GC-MS/MS was performed on flask fermentation samples, a targeted approach with quantification was used to identify particular volatile organic compounds (VOCs) of interest. Across all ScSj hybrid fermentation samples we see a varied phenotypic response (Figure 7A) with S19 producing considerably more fatty acids and their associated ethyl esters.

**Figure 7.**
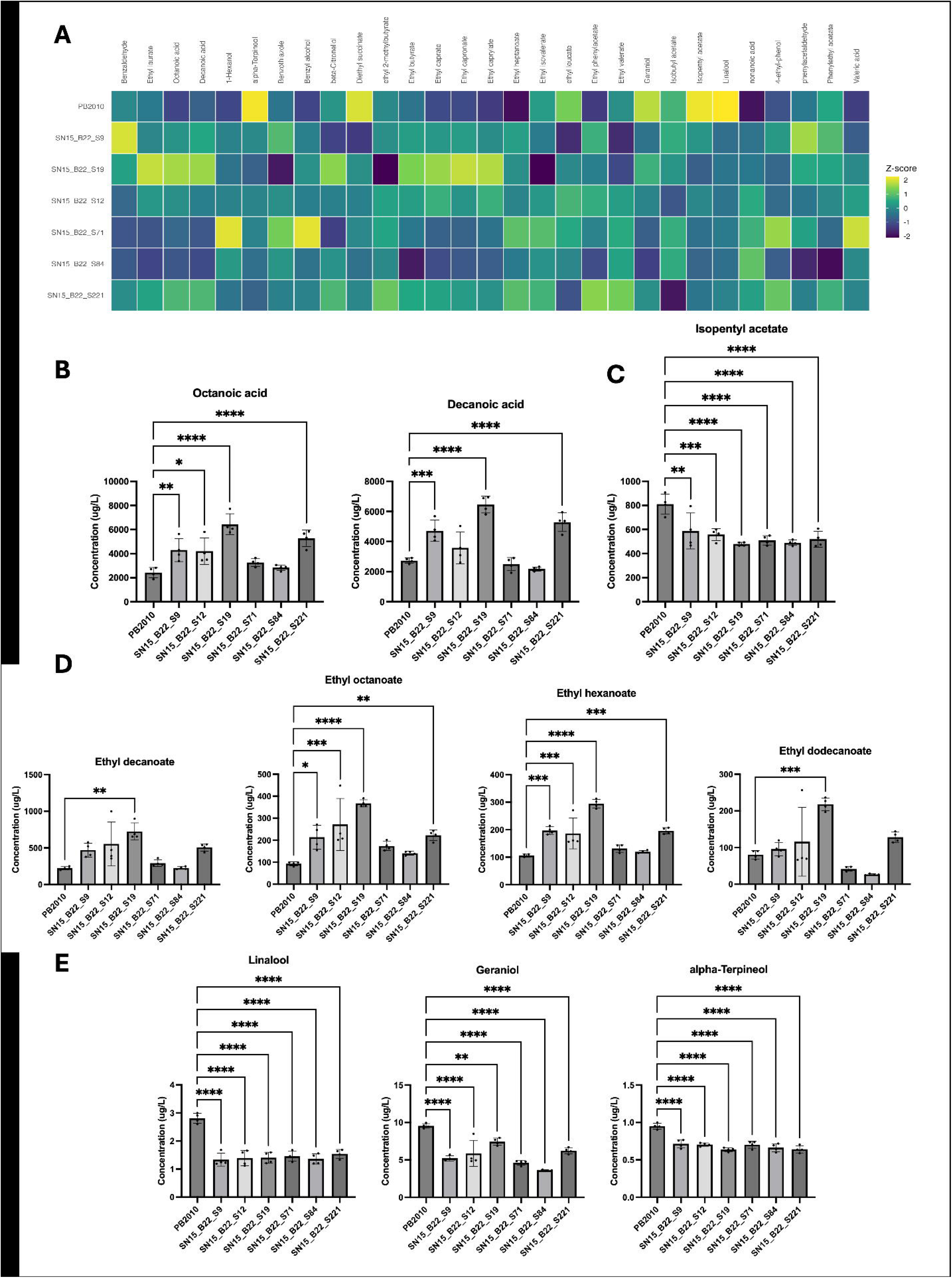
Production of VOCs detected via GC-MS/MS. **A** Comparison of production of targeted VOCs in end-point fermentation samples for hybrid progeny and PB2010 (commercial strain) in SGM media. VOCs are quantified (ug/L), heatmap displays column normalised values (z-score). **B** Quantified production of fatty acid compounds (octanoic and decanoic acids. Hybrid progeny S19 uniformly produces significantly more octanoic and decanoic acids than PB2010 and all other hybrid progeny. **C** PB2010 produces more Isopentyl acetate, a desirable ester in wine production. **D** Hybrid progeny produce greater quantities of ethyl esters than PB2010. As with fatty acid precursors, S19 uniformly outperforms PB2010 and other hybrid progeny, generating significantly more ethyl-esters. **E** PB2010 produces more terpenols. Significance across all data sets (**A**,**B**,**C** and **D**) was determined using ANOVA where; * indicate *p* < 0.05, ** indicate *p* < 0.005, *** indicate *p* < 0.0005 and **** indicates *p* < 0.0001.

We observe a higher concentration of octanoic acid in fermentations with ScSj hybrid strains (Figure 7B, Table 2). Notably, S19 produced an average of 6.441 mg/L and S221 produced an average of 5.281 mg/L of octanoic acid, a significantly higher concentration than that produced by PB2010 (2.426mg/L). Interestingly, this parallels findings in wort fermentations, with natural ScSj hybrids producing higher quantities of octanoic acid over *S. cerevisiae* strains [36]. We also observe a significantly higher concentration of decanoic acid in fermentations performed with S19 over PB2010, with concentrations of 6.451 mg/L and 2.731 mg/L respectively (Figure 7B, Table 2).

**Table 2.** Comparison of targetted quantitative GC-MS/MS data from analysis of end-point fermentation samples.

| Category | Compound | Reported aroma | Sensory threshold (ug/L) | Concentration detected (ug/L) |  |  |  |  |  |  |
|---|---|---|---|---|---|---|---|---|---|---|
|  |  |  |  | PB2010 | S9 | S12 | S19 | S71 | S84 | S221 |
| Esters | **ethyl 2-methylbutyrate** | fruity | 18 | 7.87 | 8.02 | 8.57 | 4.41 | 8.68 | 7.87 | 9.81 |
|  | **Ethyl dodecanoate** | sweet, waxy, fruity-floral | N/A | 80.54 | 95.79 | 115.92 | 218.06 | 41.47 | 26.51 | 128.11 |
|  | **Ethyl butanoate** | fruity, pineapple, sweet | 20 | **141.99\*** | **156.65** | **156.54** | **176.62** | **142.85** | **114.80** | **159.97** |
|  | **Ethyl decanoate** | sweet, apple, grape | 200 | **222.88** | **470.32** | **553.81** | **723.84** | **292.25** | **226.15** | **507.59** |
|  | **Ethyl hexanoate** | fruity, apple | 14 | **105.76** | **196.74** | **186.25** | **294.35** | **131.34** | **119.66** | **195.73** |
|  | **Ethyl octanoate** | fruity, wine, sweet | 5 | **93.13** | **213.59** | **271.73** | **367.41** | **173.37** | **140.81** | **222.33** |
|  | **Ethyl isovalerate** | fruity, apple, pineapple | 3 | **3.59** | **3.20** | **3.53** | **2.05** | **3.99** | **3.48** | **3.86** |
|  | **Ethyl leucate** | blackberry, fruity | N/A | 8.94 | 6.18 | 8.43 | 7.65 | 8.21 | 7.62 | 6.75 |
|  | **Ethyl phenylacetate** | honey, sweet, floral | N/A | 0.64 | 0.90 | 0.90 | 0.78 | 0.80 | 0.67 | 1.01 |
|  | **Ethyl pentanoate** | sweet, fruity, apple, green | N/A | 0.64 | 0.49 | 0.66 | 0.53 | 0.68 | 0.67 | 0.75 |
|  | **Diethyl succinate** | fruity, melon, berry | N/A | 13.69 | 5.66 | 8.53 | 9.15 | 8.83 | 7.78 | 8.80 |
|  | **Isobutyl acetate** | fruity, floral, banana | 1600 | 41.69 | 41.20 | 35.91 | 39.66 | 41.27 | 41.14 | 32.52 |
|  | **Isopentyl acetate** | banana, pear | 30 | 810.43 | 587.83 | 557.75 | 479.47 | 510.03 | 489.44 | 519.25 |
|  | **Phenylethyl acetate** | rose, honey, fruity | 250 | 187.15 | 196.28 | 174.78 | 186.42 | 161.97 | 128.33 | 191.83 |
| Fatty-acids | **Octanoic acid** | sweat, cheese, rancid | 500 | 2426.29 | 4305.43 | 4201.91 | 6441.47 | 3260.56 | 2835.56 | 5280.71 |
|  | **Decanoic acid** | rancid, fatty | 1000 | 2731.75 | 4712.81 | 3575.68 | 6451.10 | 2501.50 | 2177.00 | 5284.40 |
|  | **Pentanoic acid** | sweaty, cheesy, pungent | N/A | 13.85 | 14.85 | 17.19 | 16.44 | 20.33 | 16.45 | 16.28 |
|  | **Nonanoic acid** | waxy, fatty, cheesy | N/A | 34.12 | 50.23 | 61.21 | 51.32 | 62.66 | 66.48 | 59.21 |
| Aromatic alcohols | **1-Hexanol** | green, herbaceous, grassy | 8000 | 13.22 | 16.50 | 16.52 | 15.54 | 23.70 | 17.87 | 15.62 |
|  | **Benzyl alcohol** | faint, floral, sweet | N/A | 7.17 | 16.40 | 15.83 | 19.18 | 37.07 | 13.97 | 16.67 |
| Aldehydes | **Benzaldehyde** | almond, cherry | 2000 | 9.94 | 16.75 | 7.98 | 11.42 | 6.91 | 7.14 | 10.24 |
|  | **Phenylacetylaldehyde** | honey, floral, green | N/A | 15.09 | 19.22 | 14.99 | 16.97 | 16.66 | 12.43 | 15.43 |
| Terpenols and other volatile aromas | **alpha-Terpineol** | floral, pine | 250 | 0.95 | 0.71 | 0.70 | 0.64 | 0.70 | 0.66 | 0.64 |
|  | **Geraniol** | rose | 30 | 9.54 | 5.21 | 5.87 | 7.43 | 4.62 | 3.62 | 6.23 |
|  | **Linalool** | citrus, floral, waxy | 25.2 | 2.81 | 1.34 | 1.39 | 1.40 | 1.45 | 1.36 | 1.54 |
|  | **Citronellol** | rose, citrus | 100 | 8.23 | 6.99 | 7.91 | 9.39 | 6.93 | 8.19 | 8.80 |
|  | **4-ethyl-phenol** | smoky, medicinal | N/A | 0.32 | 0.99 | 0.54 | 0.51 | 1.71 | 0.42 | 1.46 |
|  | **Benzothiazole** | rubber, sulfurous | N/A | 1.63 | 1.79 | 1.69 | 1.44 | 1.85 | 1.70 | 1.65 |

*\*the numbers in bold show the concentration of volatiles above the sensory threshold*

Medium chain fatty-acids (FAs), such as octanoic and decanoic acid, can have a negative organoleptic influence at higher concentrations (>20 mg/L) in wine leading to a rancid or cheesy aroma, with a detectable limit above 0.5 mg/L and 1.0 mg/L respectively [78]. However, it has been reported that medium chain FAs can have a positive impact on wine aroma at concentrations between 4 mg/L and 10 mg/L [79, 80] with varying concentrations in wine reported [81, 82].

Esters are one of the main classes of aroma compounds in fermented alcoholic beverages, they provide fruity and wine like aroma [83-85]. Individual fatty-acid (FA) esters are not-considered to impact overall wine aroma in isolation, but a combination of medium-chain FA esters contributes to the overall quality of the product [86].

Previous analysis of non-GMO ScSj hybrids has reported an abundance of ethyl esters in beer fermentations. Specifically, when compared to *S. cerevisiae* and *S. jurei* strains in isolation, ScSj hybrids generated a higher concentration of ethyl octanoate and ethyl decanoate [36]. Our data in ScSj interspecies hybrid progeny mirror the results in Giannakou et al. 2021, with higher concentrations of ethyl esters present in end-point fermentation samples when compared with a *S. cerevisiae* commercial strain. There is a significantly higher concentration of ethyl decanoate present in S19 fermentation samples compared with PB2010 samples, an average of 0.723 mg/L and 0.223mg/L respectively. Hybrid S19 also produced significantly more ethyl octanoate (0.367 mg/L), ethyl hexanoate (0.295 mg/L) and ethyl dodecanoate (0.218 mg/L) than PB2010, as summarised in Figure 7 and Table 2. Overall, fermentations with ScSj hybrid progeny contained a greater quantity of ethyl esters in total, than the commercial strain PB2010 (Table 2). A link has been suggested between growth rate and ester production, with a higher growth rate linked to greater ethyl octanoate production [87], but also requires excess ethanol and FA precursors.

Terpene alcohols; linalool, geraniol, alpha-terpineol and beta-citronellol, were detected in end-point fermentation samples (Figure 7). Although there is a clear importance for monoterpenoids in wine aroma [88], they principally arise from grape must, prior to fermentation [89]. There is also evidence that monoterpenoids are synthesised from precursor chemicals, present in grape must, by enzymatic hydrolysis [90]. As the fermentations discussed in this paper were performed in synthetic grape must, containing no precursor chemicals, monoterpenoids present are likely due to *de novo* biosynthesis. Production of monoterpenoids has been reported in *Saccharomyces* species previously [91], at similar concentrations to those observed here. The ScSj hybrid progeny display overall lower production of terpenoids than the commercial strain PB2010 (Figure 7). With significantly higher concentrations of linalool, geraniol and alpha-terpineol present in end-point fermentation samples from PB2010 compared to each of the hybrid progeny. However, all concentrations of terpenols are lower than the sensory threshold for each detected chemical (Table 2).

### Untargeted GC×GC-ToF-MS analysis demonstrates clear separation of PB2010

We also performed an untargeted approach to identify a wider scope of aroma compounds; GC×GC-ToF-MS was performed on flask fermentation samples following previously reported strategies [36, 92]. We were able to obtain semi-quantitative data for a range of volatile compounds (VOCs), these are presented in Table S4. In total 84 compounds were identified, 14 alcohols, 24 esters and 8 acids (including fatty acids) and 37 other volatile organic compounds including aldehydes, ketones and terpenes. Principle component analysis on non-targeted chemicals shows clear clustering with the commercial strain PB2010 isolated from the hybrid progeny (Figure S4). Hybrid strains tested demonstrated broader phenotype overall, with no significant differences in grouping (Figure S4). Results observed using the targeted, quantified, approach are matched by the results obtained here (Figure S5). Hybrid S19 uniformly produces a greater amount of ethyl esters and fatty acid precursors, mirroring results obtained in targeted GC×GCMS analysis.

## Conclusions

Hybridisation offers an interesting route for development of novel flavour profiles; we use a high throughput method to investigate hybrid progeny and select promising candidates for oenological application by quantifying their production of volatile aroma compounds.

Here, we identified complex hybrid progeny between *S. cerevisiae* and *S. jurei* able to resist wine production related stressors and perform well through fermentations in complete synthetic grape must (SGM) media. Overall, we observed greater production of fatty acids and their associated ethyl esters for the Sc/Sj selected hybrids when compared with the commercial strain, PB2010. Furthermore, the undesirable production of H_2_S in these hybrids was equal or less than PB2010. In the best strain, S19, there was no decrease in the ethanol yield and higher glycerol production compared to the commercial strain. Interestingly, the hybrid S19 also outperforms all the other hybrids and PB2010 for ethyl-ester production, pivotal for the development of the wine aroma. Given the characteristic of S19, this hybrid can be a good candidate for developing new oenological strains.

Previous studies have demonstrated the clear development of interesting flavour profiles in brewing using Sc/Sj hybrids, with a smoky clove-like and spicy aroma associated with the *S. jurei* strain [35, 36].

Whilst there are limitations to wine production using genetically modified yeast strains in the EU, with debate ongoing about the use and implications of GMOs for food and beverage production [93]. However, markets exist outside the EU that are more lenient, most notably in the US and Australia [94].

## Materials and methods

### Media composition for phenotypic screening

YPD and YPDA was prepared using peptone, yeast extract, dextrose and agar. Screening was performed on SD media, consisting of YNB + ammonium sulphate (Foremedium, UK) and a complete supplementary mix of amino acid (Formedium, UK). All screening conditions included 2% glucose as standard except plates with 2% fructose, stressors were added at the following concentrations; 0.3% (v/v) and 0.5% (v/v) acetic acid, 8% (v/v) and 15% (v/v) ethanol, 200mg/L and 500mg/L potassium metabisulphite (Kmet).

### Phenotypic screening

Diploid spores of *S. cerevisiae x S. jurei* were grown in YPDA at 30°C and then inoculated into 100 µL YPD in a 96 well microtiter plate alongside the parental tetraploid (Sc^OS3^/Sj^D5088^/Sc^OS104^/Sj^D5095^). The strains were incubated for 96 hours at 30°C and then sub-cultured in 384 well microtiter plate containing 70 µL YPD using the Singer Pixl (Singer Instruments, UK), combining all 300 progeny into a single 384 layout. Liquid cultures were grown to saturation at 25°C and stamped onto YPDA solid media plates at a density of 1536 using the Singer Rotor HDA (Singer Instruments, UK), ensuring 4 technical replicates for each strain. For the phenotypic analysis of *S. cerevisiae* x *S. jurei* spores, printing onto screening plates from YPDA source plates was carried out using short pin pads (Singer Instruments, UK). The plates were imaged with the PhenoBooth Colony Counter (Singer Instruments, UK) after 72 hours of incubation. The size of the individual colonies was used as a proxy for fitness, colony size of each plate was analysed using Phenosuite software. Progeny in which a colony-size was recorded as exceeding the IQRx1.25 across the four technical replicates, were excluded from further analysis.

### Screening on BiGGY agar

Screening on BiGGY agar was performed using 384 pin pads to allow greater colony size and hence colour resolution. Images were taken on the Phenobooth (Singer Instruments, UK) and exported as raw TIF files. To assess H_2_S semi-quantitatively, ImageJ was used for analysis; plate images were converted to 8-bit greyscale and intensity (Brightness) was measured for each ROI (colonies on the plate). Greater H_2_S production results in a darker colour on BiGGY agar, this resulted in a lower intensity reading, lesser production resulted in a lighter colony size and higher intensity value.

### Synthetic grape media

Synthetic grape media (SGM) was prepared as per Viana et al. [59]. Table S5 contains all components of SGM media, ergosterol, sodium oleate, citric acid, malic acid and tartaric acid as well as sugars were added in the lab. All other components were pre-prepared as a custom order from Formedium (Foremedium, UK). 5mg sodium oleate and 15mg ergosterol were prepared in 1mL Tween80/ethanol (1:1 v/v), this was then added to 1L SGM. The pH was then adjusted to pH3.3 using 10M NaOH. SGM media used for micro-scale fermentations was prepared with all components excluding fructose and glucose, each of the stressor chemicals were added according to the listed concentrations. SGM media prepared for small-scale flask fermentations was prepared as normal with a total of 250g/L reduceable sugars (125g/L Glucose and 125g/L Fructose).

### Micro-scale fermentations

1.5mL fermentations in SGM media containing different stressors (2% (w/v) glucose, 2% (w/v) fructose, 0.3% (v/v) and 0.5% (v/v) acetic acid, 200mg/L and 500mg/L potassium metabisulphite, 12% (v/v) and 15% (v/v) ethanol, 3% malic acid, 3% tartaric acid, copper sulphate (200mg/L and 400mg/L) and at pH3.3 (adjustment using citrate-phosphate buffer [95]). Micro-fermentations were performed according to the same methods used in Giannakou et al. 2021 [36] utilising the Biolector® I system (Beckman Coulter, UK). All micro-fermentations were performed in 48-well flower plates (M2P-MTP-48-B®, Beckman Coulter, UK). Scatter-light readings with gains set to 5, 10, 20 and 30, were taken at regular intervals. Fermentation results were collected and initial analysis was performed using Biolection software (Beckman-Coulter, UK). To calculate specific growth rate, area under the curve and final expected biomass, the GrowthcurveR R package was used [96]. Robustness was calculated using the pipeline established by Trivellin et al. 2022 [40].

### Flask fermentations

Medium-scale fermentations were performed in 100mL serum bottles (Supelco, Merck) in 50mL synthetic grape media (Table S5) adapted from [59]. Fermentation bottles were sealed with crimp fitted septum lids, to ensure vials did not over-pressure, sterile 21G needles with 0.22um filters attached were inserted into lids. Anaerobic conditions were maintained by performing fermentations in an airtight 30L box (Solent Plastics, UK) fitted with an airlock, fermentation procedure is covered in greater detail in supplementary information. CO_2_ generation packs (Oxoid^TM^, Thermo Scientific) were placed in the sealed box to provide additional CO_2_ to ensure anaerobic conditions. Flask cultures were incubated at 25°C for 7 days, shaking at 150rpm. Media was collected and centrifuged to remove all cell mass and debris, cell mass and debris was collected and dehydrated in a laboratory oven set to 50°C. Supernatant was filtered through 0.22um filters and samples collected for HPLC analysis and GCxGC-MS analysis. Supernatant was stored at -80°C prior to all analysis to ensure limited loss of volatiles and to prevent further reactions from occurring.

### HPLC

Filtered supernatant was collected and 4x1mL aliquots of each fermentation sample were placed in Agilent HPLC vials with screw-caps. Metabolites were measured by HPLC on a 1260 Infinity II LC System with a Refractive Index Detector (Agilent) and a 300 × 7.8 mm Hi-Plex HPLC Column (Agilent) as described in [87]. The column was equilibrated with 5 mM H_2_SO_4_ in HPLC grade water at 55° and a 0.8 mL/min flow rate. Compounds were identified by retention time and quantified using calibration curves generated from authentic standards for Ethanol, Acetic acid, Glycerol, Glucose and Fructose.

### Quantified SPE GC-MS/MS

Samples were prepared as described previously [97], a summary is included here: the cartridge (Isolute® ENV+ Biotage) was preconditioned with 4mL dichloromethane, followed by 4 mL methanol and 4 mL model wine. 50 mL of end-point fermentation samples mixed with 100uL of internal standard (n-heptanol 250 mg/L) were loaded onto the cartridge, which was then washed with 3 mL of water. The cartridges were dried for 10 minutes, and then the free aromatic compounds were eluted with 2 mL of dichloromethane directly into the injection vials. Analyses were performed using the Agilent Intuvo 9000 system for fast GC coupled with an Agilent 7010B triple quadrupole mass spectrometer (Agilent Technologies, Santa Clara, USA) equipped with an electronic ionization source operating at 70 eV. The separation was achieved by injecting 1 μL in split mode (1:10) into a DB-Wax Ultra Inert column (30m x 0.25mm id x 0.25μm film thickness, Agilent Technology, USA). The initial temperature of the GC oven was 40 °C for 2 min, ramped up by 10 °C/min to reach 55 °C, then by 20°/min until 165°C, by 40°C/min to 240°C for 1.5 min and finally by 50°C/min to 250°C and kept at this temperature for additional 4 min (16 total runtime). Helium was used as carrier gas (with a flow of 1.2 mL/min). The mass spectra were acquired in multiple reaction monitoring mode. Nitrogen was used as the collision gas, with a flow of 1.2 mL/min in addition with Helium at 4.0 mL/min as quench gas. Compounds were quantified by the internal standard method using calibration curves prepared in model wine and processed through the same SPE procedure as the samples (R² > 0.99 for all compounds). The transfer line and source temperature were set at 250°C and 230°C, respectively. The data acquisition and subsequent analyses were done using the MassHunter Workstation software.

### Untargeted GC×GC-ToF-MS analysis

VOCs were analysed by headspace solid-phase microextraction coupled with comprehensive two-dimensional gas chromatography–mass spectrometry HS-SPME–GC×GC-TOFMS, as described previously [92]; (LECO Pegasus BTX-4D, Leco Corporation, USA). 2 mL of wine was placed in a 20 mL headspace vial, with 0.7 g of NaCl added, and 20μL of 2-octanol (2 mg/L) was used as the internal standard. Samples were equilibrated into the autosampler agitator at 35 °C water bath for 5 min, and VOCs were extracted for 30 minutes using an arrow 50/30μm DVB/CAR/PDMS (divinylbenzene/carboxen/polydimethylsiloxane) SPME fibre. The fibre was then thermally desorbed in the GC injector (250°C) in split mode (split ratio 1:25) for 2 minutes. The separation was performed with an Agilent 8890 gas chromatograph of an Agilent GC×GC–MS system. Chromatographic separation was performed using the 1st GC capillary column 30m × 0.25mm × 0.25μm film thickness VF-WAXms (Agilent Technologies), while the 2nd one was a 1.5 m × 0.15 mm ×0.15 μm Rxi 17Sil MS (Restek).

Hydrogen was used as the carrier gas at a constant flow of 1.2 mL/min. The primary oven temperature was held at 40 °C for 1.2 min, then increased at 10 °C/min to 250 °C and held for 3 min; the secondary oven and the modulator were operated at +5 °C relative to the primary oven. The modulation period was 3 s. The transfer line and ion source temperatures were set at 280 °C and 250 °C, respectively. Mass spectra were acquired in the range m/z 35-450 at an acquisition rate of 200 spectra/s, with electron ionisation at 70 eV. VOCs were identified by comparison of their mass spectra with the NIST23 mass spectral library and, where available, by comparison of linear retention indices (LRI) with previously published values. LRIs were calculated from the retention times of a series of n-alkanes (C7–C30) injected under the same chromatographic conditions, and putative identifications were retained only when the experimental LRI was within ±25 units of the reference value. Procedural blanks and pooled quality control (QC) samples were analysed throughout the sequence, injected every 10 samples. Features were removed if the ratio between the mean peak area in the samples and in the procedural blanks was lower than 10. Features whose relative standard deviation in the pooled QC samples exceeded that observed in the study samples were also discarded. Data processing was performed using ChromaTOF Sync 2D software.

### Statistical analysis

Statistical analysis was performed using GraphPad prism for robustness, analysis of growth kinetics for micro-fermentations and analysis of targeted GC-MS/MS data. Further analysis and visualisation of targeted GC-MS/MS was performed using MetaboAnalyst [98].

## Additional information

### Ethics approval and consent to participate

Not applicable.

### Consent for publication

Not applicable.

### Data Availability

All additional data is included in supplementary information.

### Author contributions

DD and WR conceived of the study. KP, RR and WR carried out phenotypic screening. WR performed micro-fermentations, flask fermentations and HPLC experiments with the input from AH. SC and FR performed GC×GC-MS experiments. All authors analysed the results. WR wrote the manuscript with the input of AH, SC, FR and DD. All authors proofread and approved the work. DD and UV provided funding.

## Acknowledgments

We wish to thank Federico Visinoni for his useful discussion and suggestions. The authors dedicate this work to the memory of Urška Vrhovšek. We are grateful for her valuable contribution to this work and wish to honour her enduring legacy as a scientist, collaborator, and friend.

